# Socioeconomic Segregation and Park Greenness: Insights Across a Strong Latitudinal Gradient

**DOI:** 10.1101/2025.03.03.640840

**Authors:** Diego Calbucheo, Horacio Samaniego

**Author notes:** Corresponding author (Horacio Samaniego). Email addresses:* (Diego Calbucheo).

## Abstract

Green Urban Areas (GUAs) in urban environments offer numerous benefits, including temperature regulation, air pollution reduction, climate event mitigation, and mental health improvement. Consequently, various organizations worldwide emphasize the importance of equitable access to quality public green infrastructure to enhance urban quality of life. Despite these efforts, global trends indicate an unequal spatial distribution of greenness and GUAs, often favoring more affluent populations. In Chile, green infrastructure availability is similarly uneven, with greener GUAs typically found in higher socioeconomic neighborhoods. Our review of the evidence reveals that this inequity exists not only between cities with different investment capacities but also within the same administrations. However, most studies focus on the Metropolitan Region and primarily consider exposure and access to greenness.

This study examines the greenness of GUAs in six Chilean conurbations, spanning a wide latitudinal gradient from Antofagasta to Puerto Montt, and compares it with socio-material segregation indicators. We calculated the mean NDVI of each green area using remote sensing data from Landsat 8. Socio-material characterization was conducted in two ways: stratifying the environment of each park using a socio-material index (ISMT) and analyzing the socioeconomic segregation of park visitors throughout the day using eXtended Call Detail Registries (XDR) from anonymized users of a major mobile telephone company (Movistar).

Our findings indicate segregation in both dimensions, with greenness inequitably distributed among different ISMT categories. Additionally, the socio-material entropy of park visitors aligns with literature trends, showing higher segregation during times when people are at home and in more peripheral and socioeconomically vulnerable areas. We discuss the need for more granular and scaled analyses and highlight the study’s relevance in addressing statistical issues such as the Modifiable Areal Unit Problem (MAUP).

## 1. Introduction

The multifaceted benefits of urban green spaces for human well-being are well documented (Iungman et al., 2023; Bai et al., 2022; Ha et al., 2022; Wicks et al., 2022). During the COVID-19 pandemic, the importance of these spaces in supporting mental health became increasingly evident (Pamukcu-Albers et al., 2021), underscoring the need for equitable distribution and accessibility across diverse socioeconomic groups (Hanzl, 2021; Wu et al., 2023). Exposure to urban green spaces is associated with numerous physical and psychological health benefits at both individual and community levels (Twohig-Bennett and Jones, 2018). Urban trees and semi-natural areas improve air quality, mitigate the effects of urban heat islands, and reduce the prevalence of mental health disorders, including anxiety and depression (Kondo et al., 2018; McDonald et al., 2021; Nutsford et al., 2013; Reyes-Riveros et al., 2021). Studies consistently demonstrate correlations between green space availability and reduced rates of respiratory diseases, neurocognitive disorders, early-onset overweight, and all-cause mortality (Ebisu et al., 2016; Islam et al., 2020; Rojas-Rueda et al., 2019). Moreover, increased access to green spaces is linked to lower levels of anxiety and depression (Astell-Burt and Feng, 2019; Dzhambov et al., 2018; Nutsford et al., 2013). Beyond human health, urban woodlands and green infrastructure provide essential ecosystem services—including biodiversity corridors and climate change mitigation—and enhance urban resilience by facilitating recovery from natural disasters (Venkataramanan et al., 2019; Eisenman et al., 2019; Escobedo et al., 2011; Kroeger et al., 2018; Bush and Doyon, 2019; Zuniga-Teran et al., 2020). Integrating green infrastructure into urban planning optimizes resource use and reduces the costs associated with climate change mitigation (O’Donnell et al., 2020; Stefanakis, 2019).

Urban development-induced habitat fragmentation necessitates the strategic planning of green infrastructure to conserve urban biodiversity (Filazzola et al., 2019; Garizábal-Carmona and Mancera-Rodríguez, 2021). Natural remnants within urban landscapes provide crucial habitats for arthropods, particularly native bees, while trees and blue-green infrastructure are vital for avian conservation (Prendergast et al., 2022; Liordos et al., 2021). Moreover, public tree management—especially in socioeconomically disadvantaged areas—has been shown to benefit urban bird populations by enhancing their presence and contributing to ecosystem services such as pest control and improved mental well-being (Wood and Esaian, 2020; Díaz and Armesto, 2003; Escobedo et al., 2011; Muñoz Varela et al., 2018). Habitat fragmentation from urban development poses a significant threat to biodiversity, highlighting the critical role of green infrastructure quality, size, and connectivity (Filazzola et al., 2019; Garizábal-Carmona and Mancera-Rodríguez, 2021).

Unequal access to urban parks is a global issue, often linked to socioeconomic factors such as income, immigration, and home ownership (Barboza et al., 2021; Csomós et al., 2020; Matthew McConnachie and Shackleton, 2010; Villaseñor and Escobar, 2022; Gerrish and Watkins, 2018; McDonald et al., 2021; Watkins and Gerrish, 2018; Johnston, 2019; Uchiyama and Kohsaka, 2022). In Chile, where nearly 80% of the population is urban (Instituto Nacional de Estadísticas, 2017), research has primarily focused on the capital city, Santiago. This focus has left a gap in our understanding of how green infrastructure evolves in larger cities and how it relates to socioeconomic segregation (Escobedo et al., 2006; Reyes Paecke and Meza, 2011; Correa Galleguillos and De la Barrera, 2014; De La Barrera and Henríquez, 2017b; De La Barrera et al., 2019). Although pioneering studies suggest that vegetation loss is concentrated in lower socioeconomic areas outside Santiago (De La Barrera and Henríquez, 2017a), further research is needed to address park access inequity—particularly in rapidly urbanizing regions—and to establish consistent “green area” definitions for accurate socioeconomic impact assessments. Chile’s current legislation lacks specificity. While existing indicators provide a quantitative measure of park quality (MINVU, 1992; CNDU, 2018), we will evaluate their relationship with overall greenness and with a specific indicator of socioeconomic segregation. Socioeconomic inequality in Chilean cities has been examined using various frameworks, yet market-based stratification remains prevalent despite its limitations (Lambiri and Vargas, 2011; Romero et al., 2012; Carrasco et al., 2022; Mena et al., 2021; Ruiz-Tagle and López M, 2014; Barozet and Espinoza, 2009; AIM, 2019). We use the socio-material index (ISMT; Índice Socio-Material Territorial) as a more holistic alternative. Segregation—measured through various metrics—remains a key indicator of inequality, with Chile exhibiting high centralization and a high Gini coefficient (Massey et al., 1996; Flores A, 2019; World Bank, 2023). Latin American cities, including those in Chile, show similar patterns of segregation, with marginalized groups becoming increasingly peripheral (Sabatini et al., 2001; Sabatini, 2006; Niembro et al., 2021; Martinez, 2015).

This study aims to characterize socioeconomic inequalities in the spatial distribution and accessibility of urban greenness across the six largest urban areas in Chile. Specifically, we will (1) analyze the spatial correlation between urban vegetation greenness and socioeconomic environments, and (2) investigate socioeconomic mixing patterns surrounding urban green spaces using a large mobile phone dataset. This approach provides a temporally resolved description of the activity spaces in these urban areas under typical, non-event conditions.

## 2. Materials and Methods

This study analyzes the six largest urban areas in Chile, which are located between 23°S and 41°S (see Fig. 1). These cities—the largest conurbations spanning different climatic regions (Di Castri et al., 1976; Flores A, 2019)—include Antofagasta, Coquimbo-La Serena, Valparaíso, Santiago, Concepción, and Puerto Montt.

**Fig. 1:**
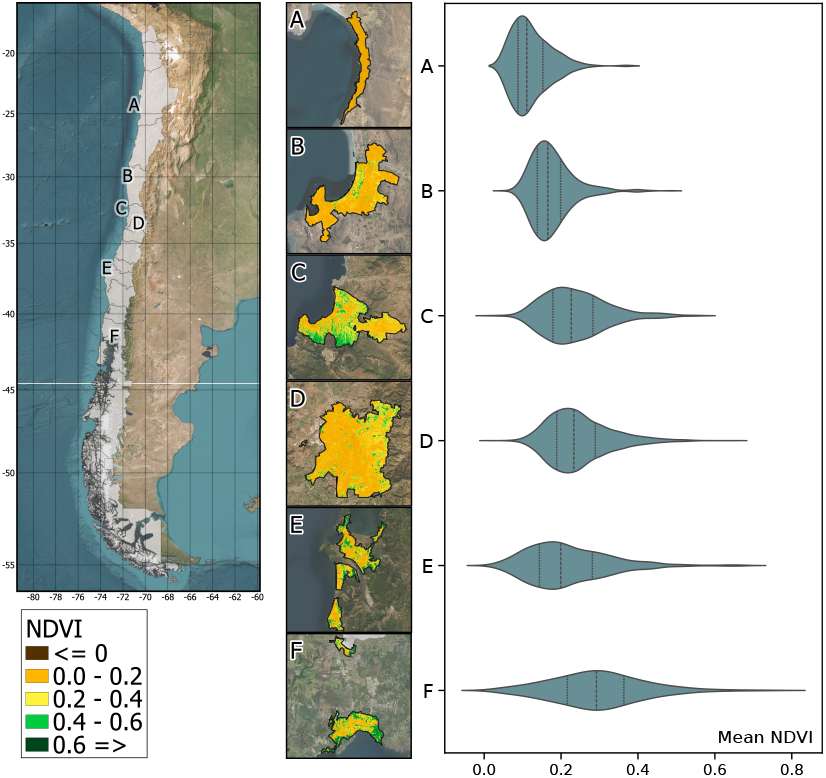
Location and greenness of the six largest conurbations analyzed in this study. Greenness is represented by the scaled values of the Normalized Vegetation Index. Violin plots in on the left represent the distribution of mean NDVI values in each city. Quartiles are shown as vertical stripes within each violin. Major conurbations are labeled by its main city: **A** Antofagasta, **B** Coquimbo-La Serena, **C** Valparaíso, **D** Santiago, **E** Concepción and **F** Puerto Montt.

We use the official database of parks and green areas developed by the Chilean Institute of Statistics, which provides data on surface area, quality, and other attributes of public green spaces across municipalities (Flores A, 2019). The quality of public parks is assessed using a Quality Index (QI) designed to rank parks based on the services they provide. This index evaluates five infrastructural components: general maintenance, type of vegetation, universal accessibility, safety, and diversity of facilities. In this study, the Quality Index is employed to analyze daily usage patterns of these public spaces.

Greenness is characterized using remote sensing imagery and quantified with the Normalized Difference Vegetation Index (NDVI), which calculates vegetation density as the ratio between the near-infrared and red bands of the electromagnetic spectrum. Higher NDVI values generally correspond to a greater fraction of photosynthetically active radiation, indicating greener, more vigorous vegetation and denser plant cover (Pettorelli et al., 2005). NDVI was computed from the Landsat 8 catalog via the Google Earth Engine platform (Gorelick et al., 2017) and is widely used to study vegetation cover in urban areas (Jenerette et al., 2011; Stefanov and Netzband, 2005). Each public park was further classified into low, medium, or high greenness categories based on its mean NDVI value relative to the citywide terciles.

Socioeconomic levels were classified using the Sociomaterial Territorial Index (ISMT), derived from the 2017 Census of Population and Housing (Observatorio de Ciudades UC, 2022). The ISMT considers the educational level of the household head, housing quality (wall, roof, and floor condition), overcrowding (persons per bedroom), and cohabitation (number of family groups per dwelling) (CASEN, 2017). Each park was assigned the ISMT value of its corresponding census zone. For parks spanning multiple zones, a weighted average ISMT was calculated based on the relative area within each zone.

To assess the activity space related to public parks, we analyzed the frequency of visits to areas surrounding these spaces. As a proxy for urban mobility, we used the eXtended Call Detail Records (XDR) from MOVISTAR, a major mobile phone provider in Chile (see Dannemann et al., 2018; Lenormand and Samaniego, 2023, for similar procedures). The dataset covers four weeks in March, May, and October 2015 and was selected to represent typical weeks without extraordinary events that might disrupt normal social and economic patterns. Each record was geolocated to a specific antenna. Like traditional Call Detail Records, XDR tracks all data activity from mobile phones connecting to nearby antennas, thereby enabling us to map individual movements throughout the city (Blondel et al., 2015).

Each city was divided into Voronoi cells to pinpoint users’ activities geographically. We then estimated the number of individuals connecting to antennas associated with public parks in one-hour intervals. Furthermore, we inferred each individual’s socioeconomic status based on their presumed residence, following the approach of Vanhoof et al. (2018). Presumed residences were defined as the most frequently visited locations between 9 PM and 7 AM, with at least five records during that period. Individuals without a determinable residence were excluded from the analysis. The data provider anonymized all records to ensure user privacy.

We categorized individuals’ socioeconomic status into four groups—low, medium low, medium high, and high—based on their ISMT quartiles within each city (Table 1). A similar procedure was applied to each park, assigning an ISMT quartile accordingly. This classification enabled us to label all public parks and individuals according to their respective socioeconomic groups.

**Table 1:** Number of users analyzed in this dataset. **Parks** represent the number of public parks considered in this study; **Area** is the total area in hectares (ha). **Antennas** is the number of antennas found in the urban area. **Q1-Q4** represent the number of individuals per ISMT quartile in increasing wealth. Population is urban population in 2015 estimated from the official census of 2012 in thousands (*×*10^3^).

| City | Parks | Area | Antennas | Q1 | Q2 | Q3 | Q4 | Population |
| --- | --- | --- | --- | --- | --- | --- | --- | --- |
| Antofagasta | 220 | 85.0 | 74 | 21485 | 10167 | 13303 | 12882 | 361.9 |
| La Serena | 996 | 311.3 | 125 | 18469 | 26624 | 21170 | 19331 | 534.2 |
| Valparaíso | 754 | 246.6 | 112 | 30693 | 34130 | 33588 | 35270 | 909.2 |
| Santiago | 9096 | 3511.4 | 42 | 323138 | 285560 | 178367 | 174442 | 6254.3 |
| Concepción | 1518 | 419.1 | 1344 | 17693 | 17237 | 12746 | 19325 | 1187.2 |
| Puerto Montt | 614 | 140.9 | 182 | 7059 | 6373 | 7385 | 6772 | 308.1 |

We estimated the number of antennas near parks by identifying the overlap between Voronoi cells and public parks. Although this method does not precisely determine whether individuals are using park facilities, it enables analysis of a large dataset that accounts for the socioeconomic stratification of individuals passing by or near green urban areas.

This approach is feasible because mobile phones connect to antennas (or “ping”) multiple times when accessing the internet (e.g., when retrieving messages), with each ping linked to the specific location of the corresponding antenna at that time.

Sankey flow diagrams were employed to qualitatively assess the association between parks’ socioeconomic status, greenness, and the Quality Index (Lupton and Allwood, 2017). Analysis of variance (ANOVA) was used to assess the effect of the classified socioeconomic status of public parks on their average greenness. The most significant differences were identified using a Tukey post-hoc test (Quinn and Keough, 2002).

We used entropy (H) as an indicator of social mixing-a measure often associated with social segregation (Massey et al., 1996; Lenormand et al., 2020; Batty, 2020). For each city, H was computed across the four socioeconomic quartiles on an hourly basis, enabling the construction of a time series that tracks changes in social mixing among public park visitors. This 24-hour time series was then analyzed for parks across different greenness terciles and ISMT quartiles. A simple Chi-square test was applied to evaluate whether socioeconomic levels were evenly distributed across greenness categories.

Finally, a cluster analysis was performed using Euclidean distances on the H time series of public parks larger than 0.5 ha. This analysis evaluated whether parks exhibiting similar temporal changes in H also shared comparable Quality Index (QI), socioeconomic status (as measured by ISMT), and greenness (NDVI). The resulting dendrograms were arbitrarily pruned to yield the four most relevant clusters.

## 3. Results

### 3.1. Greenness of Public Green Urban Areas

NDVI estimates from urban parks revealed distinct patterns in greenness distribution across the analyzed cities. Violin plots of NDVI frequency distributions indicate a latitudinal trend, with cities further south exhibiting higher average NDVI values (Fig. 1).

### 3.2. Sociomaterial Surroundings of Public Green Urban Areas

Analysis of the sociomaterial context of public green spaces revealed that in three of the six cities—namely, Antofagasta, Santiago, and Concepción—there is a direct association between ISMT and mean NDVI. This suggests that greener areas are predominantly located in higher-income neighborhoods (Fig. 2).

**Fig. 2:**
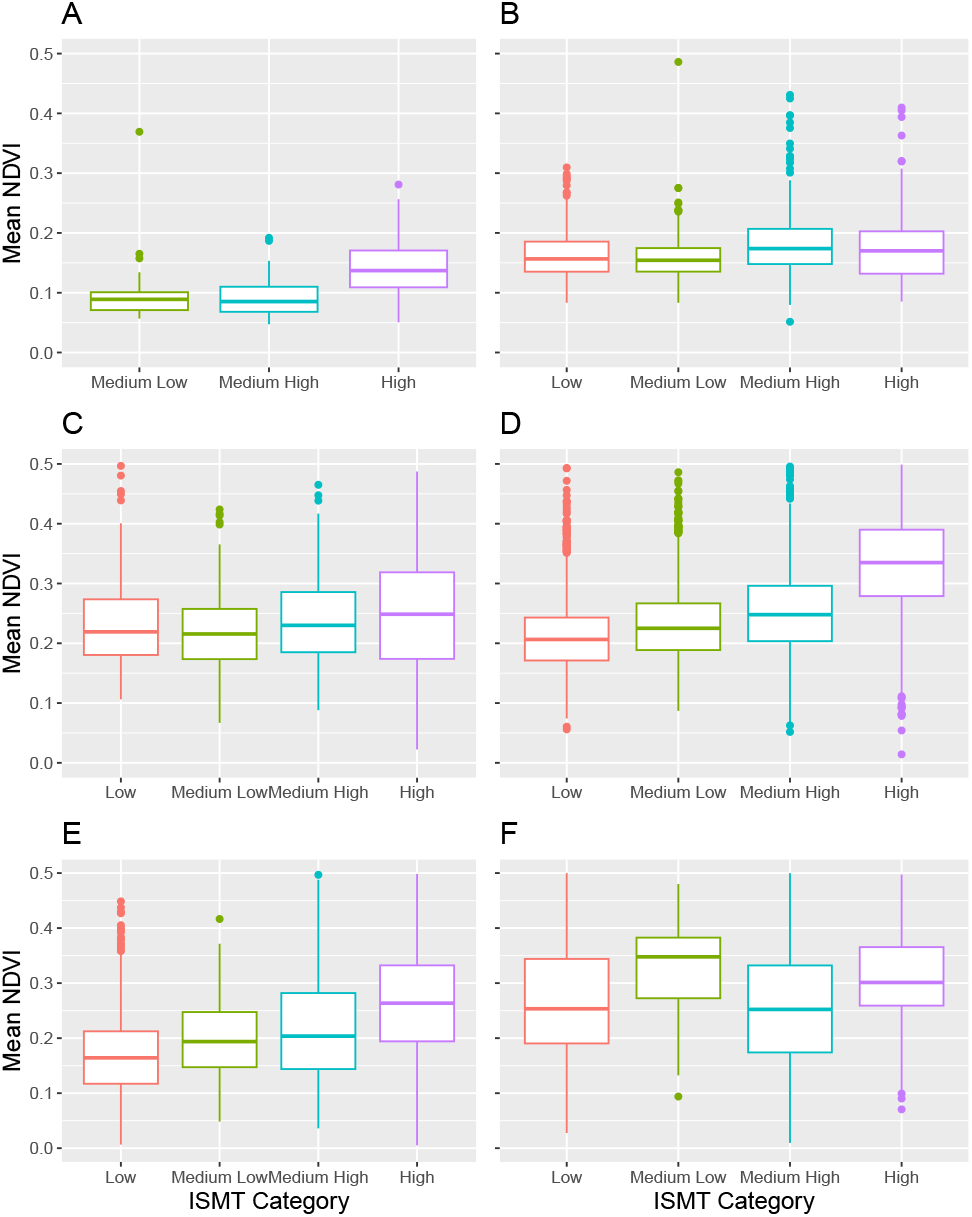
Greenness distribution among sociomaterial contexts, measured through its mean NDVI value of across six cities in Chile. The sociomaterial context of public green areas is shown for **A** Antofagasta, **B** La Serena, **C** Valparaíso, **D** Santiago, **E** Concepción, end **F** Puerto Montt. See Tukey analysis to evaluate significant differences between categories (see TableA.3).

In Antofagasta, no public parks were found in Low ISMT neighborhoods. Parks in midrange ISMT areas exhibited similar levels of greenness, with a slight upward trend, while a positive association between ISMT and greenness emerged in High ISMT areas (Fig. 2A). Although this pattern was not evident in La Serena and Valparaíso (Fig. 2B and 2C), a positive trend between ISMT and NDVI was observed in central and south central cities, such as Santiago (Fig. 2D), where higher ISMT areas consistently displayed elevated NDVI values. Notably, lower socioeconomic areas in Santiago and Concepción exhibited lower mean NDVI values, suggesting a strong link between increased greenness and higher-income neighborhoods (Figs. 2D and 2E; see Appendix A.2). In Antofagasta, although the differences in greenness between Medium-Low and Medium-High ISMT areas were small, the Tukey test revealed significant differences in mean NDVI values between High and Medium-Low income areas (*p <* 10^−4^, 95% C.I. = [0.025, 0.064]) and High and Medium-High income areas (*p <* 10^−4^, 95% C.I. = [0.035, 0.068]). In Santiago, the largest mean NDVI difference (Δ) was observed between High and Low ISMT levels (Table A.3). Despite an overall upward trend, the differences between Low and Medium-Low, and between Medium-Low and Medium-High levels, were relatively small (Fig. 2D with ΔNDVI = 0.020 and 0.021, respectively, see Table A.3). A similar pattern was observed in Concepción, where areas in the Low ISMT category exhibited the lowest greenness (Fig. 2E). The largest ΔNDVI was found between the extreme ISMT categories, with the next largest difference occurring between Medium-High and High ISMT categories, mirroring the trend seen in Santiago (Table A.3). Although the cities of La Serena, Valparaíso, and Puerto Montt did not show an evident association between ISMT level and mean NDVI, in all cases, the ANOVA revealed a significant difference between mean NDVI among ISMT levels (Table A.2).

The Sankey flow diagram in Fig. 3 shows that in Antofagasta, 88% of parks with High greenness are in High ISMT areas, while only four and five parks are in Medium-Low and Medium-High ISMT areas, respectively. Figure 3A shows that few parks are in Medium-Low income ISMT areas, with half in High ISMT areas (i.e. 128 out of 220). In La Serena, the greenness distribution among ISMT levels is less clear, but 67% of parks are in higher ISMT groups, and only 18% are in lower ISMT areas (Fig. 3B). Valparaíso shows a different pattern, with the fewest parks in the lowest and highest ISMT areas, having 152 and 125 parks out of 760, respectively (Fig. 3C). In Santiago, the Medium-High and High ISMT areas have the fewest parks, with 2,034 and 1,476 respectively out of 9096 total parks (Fig. 3). Conversely, Low and Medium-Low ISMT areas host the majority of public parks with 3,343 and 2,638 parks respectively (i.e. a 65.75%). Despite this, 80% of parks in high ISMT areas are categorized as High greenness, while this proportion decreases with ISMT. For example, 48% of parks in Low ISMT areas are of Low greenness, compared to 8% in High ISMT areas (Fig. 3D). In Concepción, like Santiago, most parks (i.e. 584) are in lower ISMT groups, with 48% showing low greenness. This proportion decreases as ISMT levels increase. Conversely, parks with high greenness are underrepresented in low ISMT quartiles (i.e. 15% and 24%), while those in high ISMT levels constitute a nearly 60% of the total (Fig. 3E). In Puerto Montt, low ISMT areas have significantly fewer parks (54 and 88, respectively) compared to higher ISMT levels, with 279 and 194 parks, respectively (Fig. 3F).

**Fig. 3:**
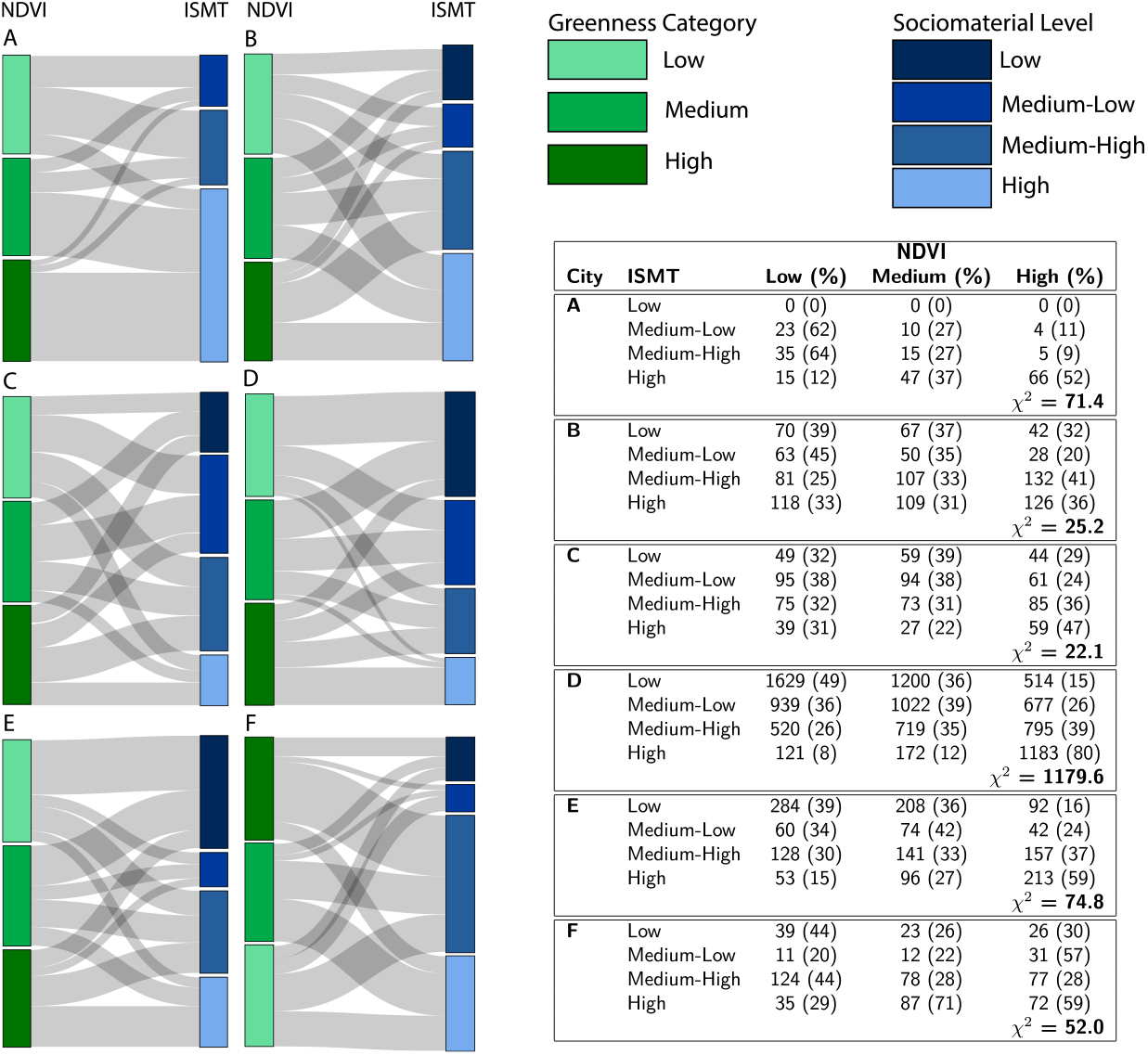
Sankey flow diagrams showing the relationship between socioeconomic condition and greenness level in **A** Antofagasta, **B** La Serena, **C** Valparaíso, **D** Santiago, **E** Concepción, end **F** Puerto Montt. Socioeconomic conditions is measured by ISMT in quartiles and greenness is the mean NDVI for each public park categorized in terciles. Frequency table shows the number of parks in each ISMT quartile distributed among greenness categories. *χ*^2^tests were computed with *df* = 4, and *p* − *value <* 10^−5^for all six tests. The percentage of parks in each category is in parentheses.

*χ*-square tests show a statistically significant (p-values *<* 2 *×* 10^−4^) disparity in the distribution of parks among each of the NDVI and ISMT categories for all cities.

### 3.3. Segregation of Public Green Urban Area Users

The hourly variation of average entropy (H) highlights differences in social mixing patterns across park visits in various cities (Fig. 4). While cities generally exhibit similar H trends—lower values at night and higher during working hours—distinct patterns emerge based on park characteristics such as greenness or ISMT. In the northern cities of Antofagasta, La Serena, Valparaíso, and Santiago, parks in higher socioeconomic areas consistently show elevated H values, indicating greater social mixing throughout the day (Fig. 4A-D). Similarly, parks with higher greenness exhibit higher H values. In contrast, the southern city of Concepción demonstrates lower H values in parks located in areas with higher ISMT levels, with greater variability and less consistency in H trends based on greenness or ISMT (Fig. 4E).

**Fig. 4:**
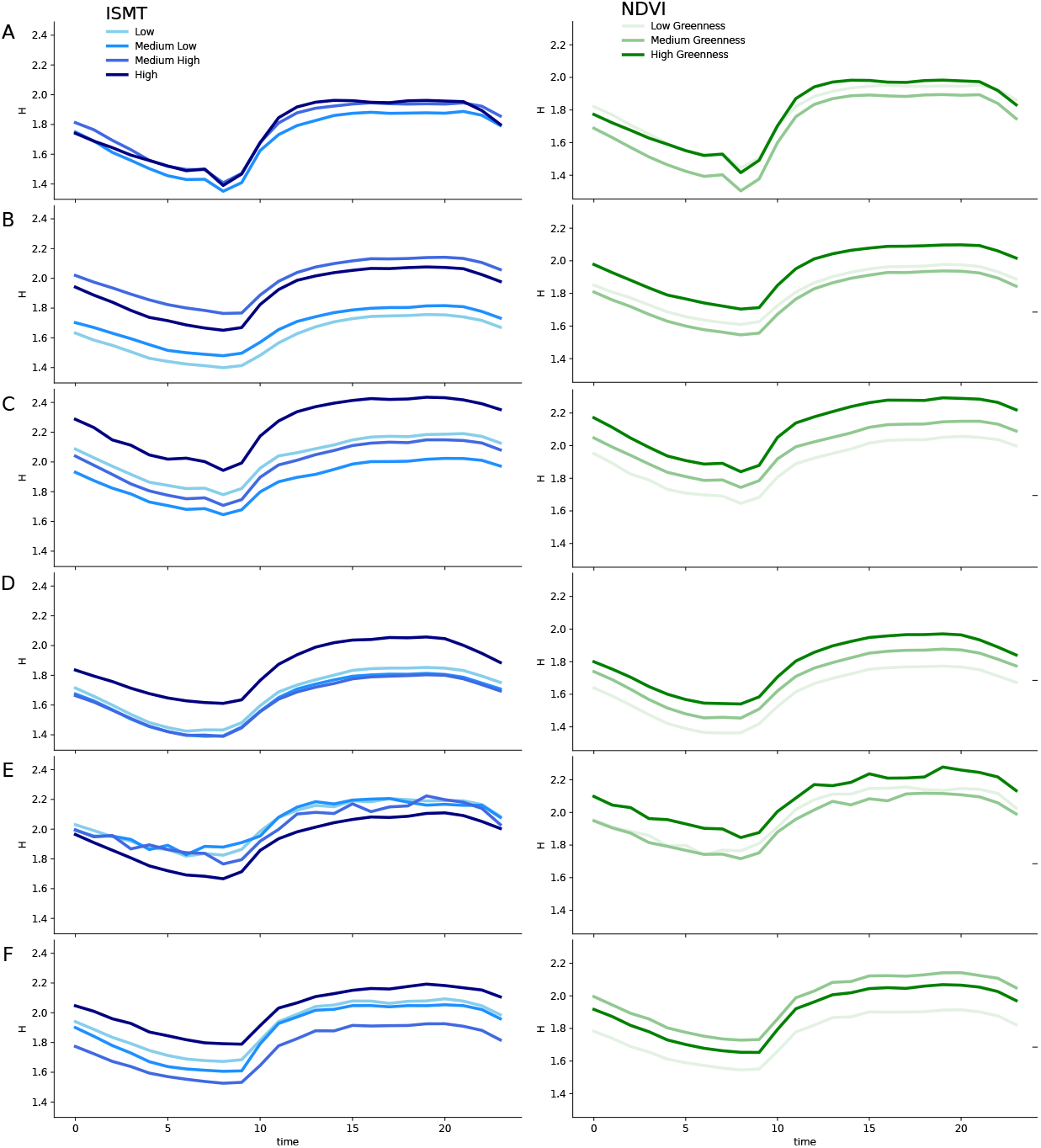
Timeline plots showing changes in entropy (H) across the day plotted across ISMT and NDVI quartiles and terciles, respectively. ISMT quartiles are shown in a gradient of blues and mean NDVI terciles in green. Cities are in rows: **A** is Antofagasta, **B** La Serena, **C** Valparaíso, **D** Santiago, **E** Concepción, and **F** Puerto Montt. Socioeconomic conditions is measured by ISMT in quartiles in a gradient of blues, and greenness is mean NDVI in greens for each public park categorized in terciles.

Clustering analysis revealed mixed associations (Fig. 5). Except for Antofagasta, no consistent relationship was identified between QI, NDVI, or ISMT and cities with similar H patterns. In Antofagasta, public parks in clusters (iv) were associated with lower NDVI and ISMT levels, whereas clusters (i) and (ii) seem to be associated to greener and more affluent areas (i.e., higher NDVI and ISMT). Detailed dendrograms and cluster compositions are provided in the Appendix.

**Fig. 5:**
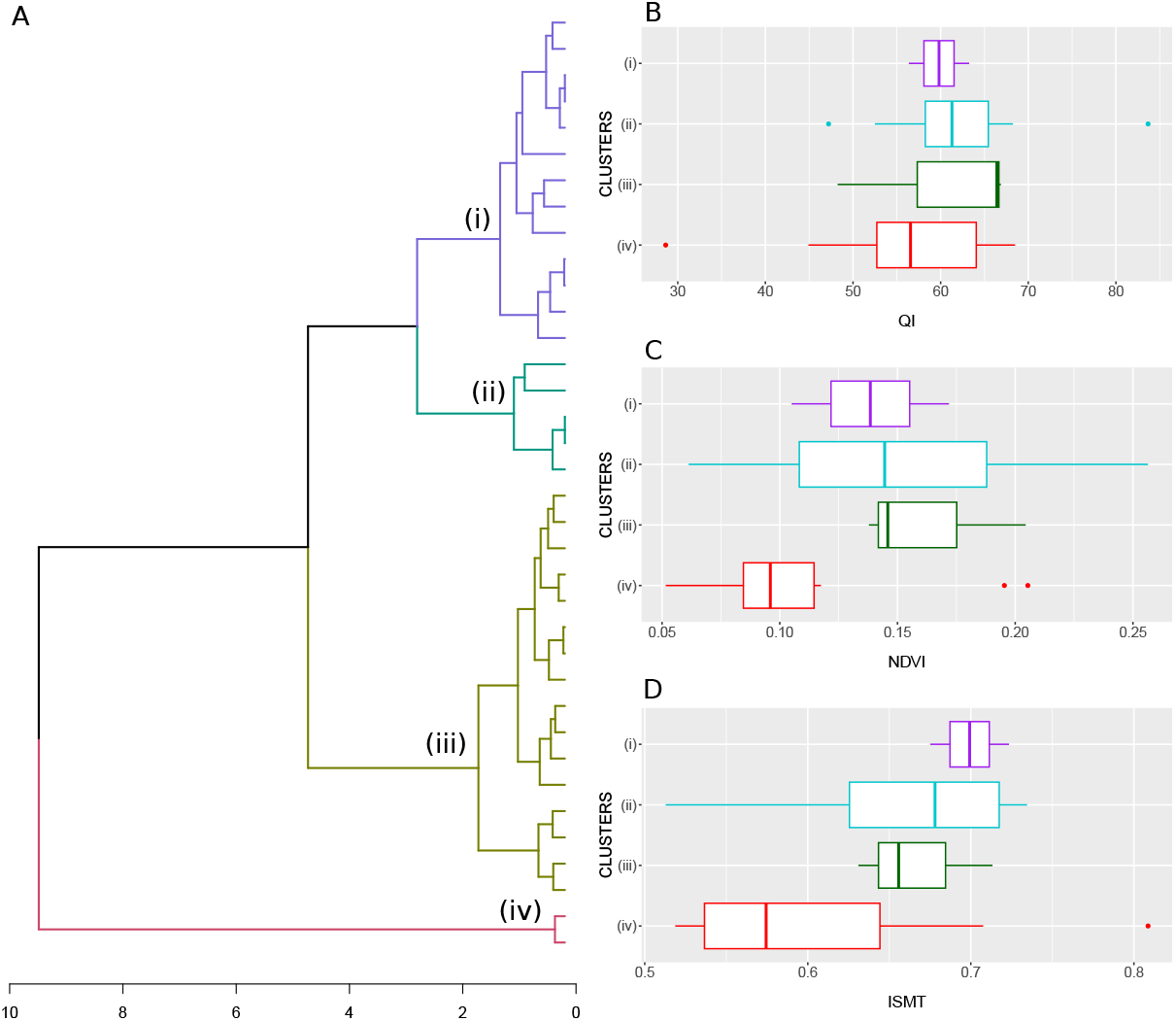
Cluster analysis of the entropy timelines of public parks for Antofagasta. **A** shows the dendrogram arbitrarily trimmed to four clusters, *i.e*. (i) to (iv) represent parks sharing a similar patterns of entropy timelines across the day. Values on the *x-axis* show the distance between clusters evaluated using the UPMGA algorithm. **B** shows the boxplot of the Quality Index of public parks among clusters in **A. C** shows the differences between mean NDVI values across clusters. **D** the mean ISMT distributed across clusters.

## 4. Discussion

Our study reveals a contrasting trend in park visitation across Chile’s strong latitudinal gradient (23^°^38’S to 41^°^28’S). Despite wealthier areas tend to have greener parks, they exhibit lower segregation levels compared to poorer areas. This pattern remains consistent across diverse climatic regions, suggesting that the unequal distribution of urban greenness is influenced more by political and administrative decisions rather than solely by climate. For example, disparities in municipal investments for park maintenance have been documented in Santiago (Escobedo et al., 2006), and our findings suggest similar patterns across other major Chilean cities, reinforcing evidence of socioeconomic segregation in urban green space exposure (Méndez et al., 2021).

The relationship between park greenness and socioeconomic status aligns with previous research (Nesbitt and Meitner, 2016; Fossa et al., 2023; Klompmaker et al., 2023), spanning from the Atacama Desert (23.5^°^S) to the Valdivian rainforest (36.8^°^S). However, in some cities—such as Coquimbo-La Serena, Valparaíso, and Puerto Montt—this relationship is less pronounced, likely due to local urban structure, neighborhood-scale variations, and planning policies (Engelberg et al., 2016; de Vries et al., 2020). Determining whether greenness disparities are primarily shaped by socioeconomic status or by other variables such as population density, latitude, or education levels remains a challenge (Wu et al., 2022; Viinikka et al., 2023).

Hourly visitation entropy patterns were consistent across cities, with higher social heterogeneity during working hours (7:00-19:00) and greater segregation during non-working hours (20:00-6:00). These findings reinforce previous studies on temporal segregation dynamics in urban spaces (Zhang et al., 2022). Concepciónexhibited unique entropy fluctuations, which may be influenced by its urban density, diversity of services, or commuting patterns (Cai et al., 2024).

A key limitation of this study is the imprecision of mobile phone-based park visitation data. Because users are assigned to antenna coverage areas rather than exact park locations, some non-visitors may be included in the dataset. While this issue is mitigated by the large sample size, integrating on-site surveys or high-resolution GPS data would improve the accuracy of visitation estimates, specially for smaller parks.

Additionally, our analysis is affected by the Modifiable Areal Unit Problem (MAUP) (Fisher and Langford, 1995; Garreton and Sánchez, 2016) which influences the granularity of entropy measurements. In particular, smaller parks that fall within a single antenna may have less precise segregation estimates. Adjusting the spatial resolution of analysis and integrating alternative geospatial thecniques could help refine these assessments (Garreton et al., 2020).

Despite these challenges, this study highlights the value of large-scale mobile phone data in exploring socioeconomic segregation in urban green space use. By identifying disparities in park greenness and visitation patterns, our findings provide actionable insights for urban planning and policy development. Future research should incorporate alternative datasets, improved spatial methodologies, and multi-scale analysis to deepen understanding and strengthen policy recommendations (Candipan et al., 2021; Kephart, 2022).

## Acknowledgements

HS thanks the chilean national agency for research and developments through its FONDECYT Regular grant #1211490.

## Appendix A. Supplementary Material

### Appendix A.1. Analysis of Variance

The ANOVA helps to determins the statistical differences between the average greenness associated to socioeconomic quintiles of green urban areas.

**Table A.2:** ANOVA table showcases the significance of the differences between average socioeconomic quintiles among the cities analyzed. Note that all cities show significant differences of average NDVI (greenness) between socioeconomic (ISMT) quintiles.

|  | DF |  | Sum Sq |  | Mean Sq |  | F Value | p-value |
| --- | --- | --- | --- | --- | --- | --- | --- | --- |
|  | Q | Residuals | Q | Residuals | Q | Residuals |  |  |
| **A** - Antofagasta | 2 | 217 | 0.129 | 0.419 | 0.065 | 0.002 | 33.433 | $2 \times 10^{-13}$ |
| **B** - La Serena | 3 | 998 | 0.069 | 2.757 | 0.023 | 0.003 | 8.410 | $10^{-5}$ |
| **C** - Valparaíso | 3 | 761 | 0.140 | 4.860 | 0.047 | 0.006 | 7.336 | $7 \times 10^{-5}$ |
| **D** - Santiago | 3 | 9464 | 16.13 | 44.522 | 5.377 | 0.005 | 1143 | $< 2 \times 10^{-16}$ |
| **E** - Concepción | 3 | 1534 | 2.1767 | 14.290 | 0.725 | 0.009 | 77.886 | $< 2 \times 10^{-16}$ |
| **F** - Puerto Montt | 3 | 607 | 0.363 | 8.789 | 0.121 | 0.015 | 8.352 | $< 2 \times 10^{-5}$ |

### Appendix A.2. Tukey tests

Tukey tests determines the specific quintiles exhibiting statistical significant differences of average greenness between green urban areas stratified by its socioeconomic quintile.

### Appendix A.3. Dendrograms

We here show dendrograms for the five remaining cities, *i.e*. La Serena, Valparaíso, Santiago, Concepción and Puerto Montt.

**Table A.3:** Tukey post-hoc analysis shows the mean NDVI differences between ISMT quartiles. Significant comparisons between groups I and J are highlighted and marked by an asterisk (*i.e. p <* 0.05).

| City | (I) ISMT Quartile | (J) ISMT Quartile | Mean difference<br>(I - J) $\Delta$ NDVI | 95% Confidence Interval<br>(lower, upper) |
| --- | --- | --- | --- | --- |
| **A** - Antofagasta | Medium-Low | Medium-High | 0.007 | (-0.015, 0.029) |
|  |  | **High** | **-0.045\*** | 0.064, 0.025) |
|  | Medium-High | **High** | **-0.052\*** | (0.068, 0.0350) |
| **B** - La Serena | Low | Medium-Low | -0.003 | (-0.018, 0.013) |
|  |  | **Medium-High** | **0.019\*** | (0.007, 0.032) |
|  |  | High | 0.011 | (-0.024, 0.001) |
|  | Medium-Low | **Medium-High** | **0.022\*** | (-0.036, -0.008) |
|  |  | **High** | **-0.014\*** | (-0.027, -0.001) |
|  | Medium-High | High | 0.008 | (-0.002, 0.018) |
| **C** - Valparaíso | Low | Medium-Low | -0.015 | (-0.036, 0.006) |
|  |  | Medium-High | 0.005 | (-0.017, 0.026) |
|  |  | **High** | **-0.025\*** | (-0.050, -0.001) |
|  | Medium-Low | **Medium-High** | **-0.019\*** | -0.038, -0.001) |
|  |  | **High** | **-0.040\*** | (-0.063, -0.018) |
|  | Medium-High | High | -0.021 | (-0.063, 0.002) |
| **D** - Santiago | Low | **Medium-Low** | **0.020\*** | (0.016, 0.025) |
|  |  | **Medium-High** | **0.042\*** | (0.037, 0.047) |
|  |  | **High** | **-0.123\*** | (-0.128, -0.117) |
|  | Medium-Low | **Medium-High** | **-0.021\*** | (-0.026, -0.016) |
|  |  | **High** | **-0.102\*** | (-0.108, -0.097) |
|  | Medium-High | **High** | **-0.081\*** | (-0.087, -0.075) |
| **E** - Concepción | Low | **Medium-Low** | **0.024\*** | (0.002, 0.045) |
|  |  | **Medium-High** | **0.048\*** | (0.032, 0.064) |
|  |  | **High** | **-0.097\*** | (-0.114, -0.081) |
|  | Medium-Low | **Medium-High** | **-0.024\*** | (-0.046, -0.002) |
|  |  | **High** | **-0.074\*** | (-0.096, -0.051) |
|  | Medium-High | **High** | **-0.050\*** | (-0.067, -0.032) |
| **F** - Puerto Montt | Low | **Medium-Low** | **0.058\*** | (0.004, 0.112) |
|  |  | Medium-High | 0.003 | (-0.035, 0.041) |
|  |  | **High** | **-0.050\*** | (-0.090, -0.010) |
|  | Medium-Low | **Medium-High** | **0.055\*** | (0.009, 0.101) |
|  |  | High | 0.008 | (-0.040, 0.056) |
|  | Medium-High | **High** | **-0.047\*** | (-0.076, -0.018) |

**Fig. A.6:**
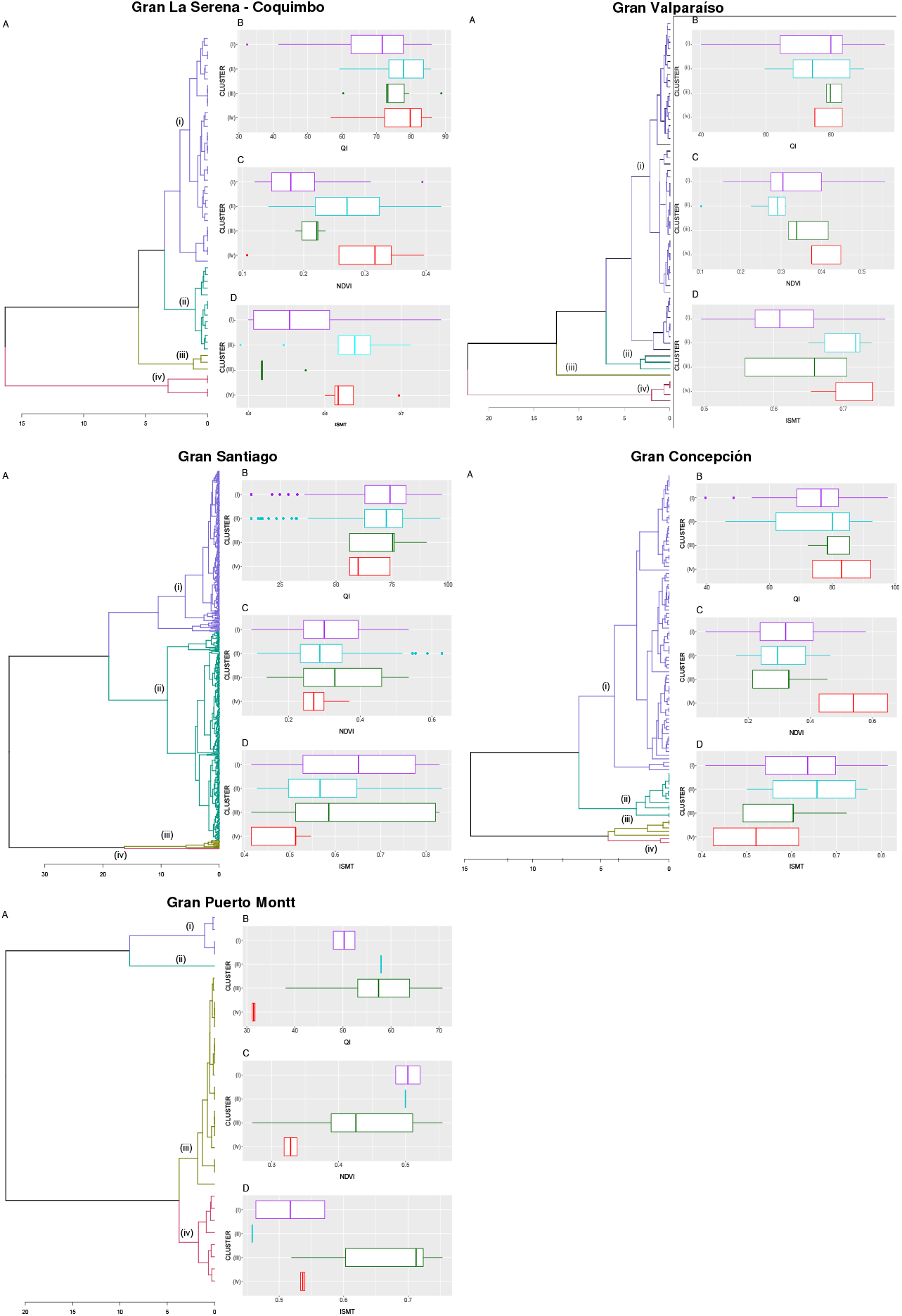
Cluster analysis of the entropy timelines of public parks for La Serena, Valparaíso, Snatiago, Concepción and Puerto Montt. **A** shows the dendrogram arbitrarily trimmed to four clusters, *i.e*. (i) to (iv) represent parks sharing a similar patterns of entropy timelines across the day. Values on the *x-axis* show the distance between clusters evaluated using the UPMGA algorithm. **B** shows the boxplot of the Quality Index of public parks among clusters in **A. C** shows the differences between mean NDVI values across clusters. **D** the mean ISMT distributed across clusters. note that Antofagasta is shown in the main text of the manuscript.

